# Persistent Cognitive Impairment and Bone Health Following Helium (^4^He) Ion Exposure: Implications for Space Travel

**DOI:** 10.64898/2025.12.08.692986

**Authors:** Laura C. Bowman, Victoria E. Elliott, Sofia Manicka, Julia Schaffer, Rosalie R. Connell, Patricia K. Thomas, Calvin J. Okulicz, Joan Smith, Alexis Mraz, Anthony G. Lau, Catherine M. Davis

## Abstract

The current study investigated the long-term neurobehavioral and physiological effects of low-dose helium (^4^He; 250 MeV/n) ion exposure, a significant component of galactic cosmic radiation (GCR), on male Long Evans rats. Groups of rats were trained on the rodent psychomotor vigilance test (rPVT) and irradiated at the NASA Space Radiation Laboratory at Brookhaven National Laboratory. Following exposure, they were tested in the rPVT from 30 – 180 days, and assessed for social recognition memory at 30, 90, and 180 days. At each time point, blood and bone samples were collected for subsequent analysis. We found that acute exposure to ^4^He ions (25 cGy) significantly impaired sustained attention, resulting in increased lapses in attention, increased reaction time measures, and decreased correct responses. However, exposure to 5 cGy ^4^He ions only increased some reaction time measures in the rPVT. Both groups displayed impaired social recognition memory. When deficits were found, they persisted for 180 days post-exposure, showing no signs of recovery. Notably, impairments in sustained attention appeared to worsen over time. While these low exposure doses of ^4^He did not significantly alter overall bone mechanical properties, specific individual parameters of bone strength were affected, and age-related changes likely played a role in observed skeletal integrity changes. Furthermore, we identified significant correlations between circulating cytokines (TNF-α, IL-1β), undercarboxylated osteocalcin (ucOC) levels, and bone biomechanical properties with behavioral performances. These findings suggest the potential for these blood- and bone-based targets to serve as diagnostic biomarkers for radiation-induced neurobehavioral effects, opening avenues for future countermeasure development. The sustained and progressive nature of the neurobehavioral deficits observed underscores the critical need for effective countermeasures to protect astronaut health and performance during exploration-class missions.

## Introduction

Future long-duration exploration class missions will involve travel outside of the protection of Earth’s magnetosphere, exposing astronauts to galactic cosmic rays (GCR) comprised of high energy and charge (HZE) ions^1^. Ionizing radiation exposures acquired on Earth, primarily through radiotherapy, support the radiosensitivity of numerous body organs, including the central nervous system (CNS) and bone^2–4^. However, GCR is qualitatively different than low linear energy transfer photon radiation, with astronauts being exposed to different ions, doses, and dose rates than those used in most radiotherapy exposures. Importantly, it is currently unknown how space radiation exposure might negatively impact astronaut cognitive performance and skeletal properties that could impair operational task performance during long-duration missions. A greater understanding of these potential radiation-induced changes is needed to avoid potentially catastrophic changes in cognition and/or increased risk of fracture.

Numerous rodent studies have independently reported simulated space radiation-induced damage to the CNS and deleterious changes in bone. For the CNS, these changes include significant impairments in recognition and spatial memory ^5–9^, social behavior ^5,10–13^, cognitive flexibility ^14,15^, and anxiety-like behaviors, in addition to increased oxidative stress^9,16,17^, autophagy^18^, microglial activation^19^, reduced dendritic complexity^20,21^, and changes in neuronal function^12,22^. Bone, a critical organ for various body functions, is similarly affected by radiation exposure. For example, high dose radiation causes a loss in cancellous bone volume and total volume, suggested to be caused by impairment to osteoblast genesis^23^. Further, ionizing radiation exposure damages collagen and hydroxyapatite, which ultimately decreases bone ratio density and stiffness^24^. Bones exposed to simulated space radiation show chronic suppression of bone formation, increases in redox-related gene expression, and trabecular bone loss ^25–27^. While the CNS and bone are often thought to function independently, recent work has shown important interactions between these organs, including bone hormones that can cross the blood-brain barrier and affect the structure and function of the CNS, in addition to links between skeletal integrity and cognitive function^28–32^. Few, if any, studies have assessed the effects of space radiation exposure on both systems in the same subjects, which would provide a method to understand potential links between radiation-induced changes in the brain and skeletal system, and advance novel mechanisms for biomarker and countermeasure development.

Osteocalcin, a hormone secreted exclusively by osteoblasts, is a non-collagenous protein that is present in two forms, carboxylated (cOC) and undercarboxylated (ucOC), with cOC an important inhibitor of bone mineralization^32^, and ucOC found to have essential extra-skeleton functions, including regulating metabolism, insulin secretion, and modulating neurotransmitter synthesis and cognition^28,33^. Interestingly, peripheral ucOC levels have been correlated with cognitive and metabolic outcomes in both humans and animals^29,31,34–40^. However, no studies to date have examined the potential interaction of radiation exposure on the CNS, bone, and osteocalcin.

In the current study, we sought to determine the effects of acute helium (^4^He, 250 MeV/n) ion exposure on the CNS and bone in the same animals, such that any potential relationship between CNS and bone radiation outcomes could be explored. Helium ions were used because they are one of the most abundant ions in GCR, but relatively little is known about their biological effects. Male Long Evans rats were irradiated with one of two doses of ^4^He (5 or 25 cGy), tested in several behavior assays following radiation exposure, and evaluated for bone mechanical properties and blood-based targets (e.g., cytokines, hormones) associated with radiation exposure.

## Materials and Methods

### Subjects and Apparatus

Laboratory animal care was conducted according to Public Health Service (PHS) Policy on the Humane Care and Use of Laboratory Animals, and the Institutional Animal Care and Use Committee of the Johns Hopkins University (JHU), Uniformed University of the Health Sciences (USUHS), and Brookhaven National Laboratories approved all procedures. JHU and USUHS also maintain accreditation of their programs by the Association for the Assessment and Accreditation of Laboratory Animal Care (AAALAC). Male Long-Evans rats (N=83, Envigo, East Millstone, NJ) were received in the laboratory at approximately 10–12 weeks of age and irradiated at approximately 5-6 months of age. Rats were singly housed in individual plastic cages and maintained on a 12:12 h light-dark schedule (lights on at 0600) and at an ambient temperature of 23°C for the duration of the experiment. Rats were run in 30-min rPVT sessions at the same time each day in identically constructed operant chambers. Each rPVT chamber contained one nose-poke key, cue lights, a house light, and a food cup for delivery of food pellets. All chambers were contained in sound-attenuating enclosures equipped with an exhaust fan. The weights of the rats were maintained at 90% of each rat’s free-feeding weight by feeding measured amounts of rat chow each day (30 min after the experimental session, 5 days/week; at similar times on the weekends), in addition to the food that was earned during the behavioral test sessions. Water was freely available in the home cage. For the rPVT procedure, experimental contingencies were controlled by MedPC V (Med Associates, Burlington, VT) behavioral control programs running on PCs; the programs recorded all data on a trial-by-trial basis to provide for a wide range of subsequent analyses. During the three-day Social Order Recognition Memory (SORM) test, no other behavioral testing was conducted. All rats had continuous access to enrichment toys (wooden blocks (K3511), Chewy Nylabones (K3584) and polycarbonate tunnels (K3325) (all from BioServ, Flemington, NJ) when they arrived in the laboratory and for the duration of the experiment. All behavioral testing was performed in the light phase between 0800 – 1530.

### Rodent Psychomotor Vigilance Test (rPVT)

Rats were first trained to respond on a nose-poke key for food pellets on a fixed-ratio 1 schedule of reinforcement. Once this behavior was acquired, training on the rPVT procedure then began. Sessions began with the onset of the house light. After a variable delay of 3–10 sec the light behind the nose-poke key was illuminated. A correct response was defined as a response on the nose poke key within 1.5 sec after the light onset (i.e., 1.5-sec limited hold, LH) and was reinforced with a pellet. A response prior to the light onset (premature response) was not reinforced and punished with an 8 sec time out, while a response after the 1.5-sec interval had elapsed (omission) was not reinforced. The delay period for the next trial began after a 1-sec inter-trial interval, timed either after the response or the end of the 1.5-sec LH, whichever occurred first. Data collected were the numbers of correct responses as defined above, premature responses (responses that occurred prior to the light onset), omissions (number of trials with no response), and lapses in responding (omissions plus responses greater than twice each rat’s mean response latency for that session). Summary measures were expressed as percentages, as follows: accuracy = number of correct responses / (corrects + premature responses + omissions); premature responding = number of premature responses / (corrects + premature responses + omissions); lapse rate = (omissions + lapses) / (corrects + premature responses + omissions). In the human literature on PVT performance, lapses are considered an important indicator of inattention and/or fatigue and are typically defined as responses with latencies greater than 500 msec, or roughly twice the average latency for humans performing the 10-min version of the test. For rodents, average latency can vary considerably from subject to subject, and so the definition adopted here was based on each rat’s individual mean latency measure which were previously described in detail^41^. Premature responding was broken down further by calculating a false alarm (FA) rate for those premature responses occurring within the 3-10 second delay period (i.e., FA rate = premature responses within the 3-10 sec delay interval / (corrects + premature responses within the 3-10 sec delay interval + omissions). Percent correct (PC) scores and false alarm (FA) rates were converted into z scores, and subtracted d’ = z(PC)-z(FA)^42^. These performance measures are important for calculating a d prime (*d’*) measure of discriminability that can be compared across the different animals. To acquire a single value that described daily rPVT performance for each subject, we computed an overall performance score that is similar to the overall performance score used in the human PVT literature^43,44^: Overall Performance Score (OPS) = 1 - (Premature Responses + Lapses)/Total Trials (including Premature Responses). The OPS ranges from 1 (perfect performance) to 0 (worst possible performance). Finally, response latencies to the light onset were recorded in milliseconds, and summarized by calculating various reaction time distributions (e.g., 10^th^ percentile, median, 90^th^ percentile, mean).

The criterion for inclusion in the present study was that rats achieve at least 75% response accuracy and less than a 25% false alarm rate, resulting in a *d’* index of 1.35, for four out of the five daily test sessions during each of the two weeks prior to radiation. In order to assign rats to radiation dose groups, a pseudo-random ranking technique was used in a manner identical to our previous work^41,45–47^. Specifically, the *d’* index for each rat was used and rats were ranked from highest to lowest *d’* and were then assigned to one of the three dose groups (0, 5, or 25 cGy) such that the average *d’* index was comparable across groups. The average pre-exposure *d’* index for each group was: sham control = 2.61 (±0.64 SD), 5 cGy = 2.69 (± 0.53 SD), and 25 cGy = 2.69, (± 0.55 SD). Following radiation exposure, rats were tested daily (M-F) in the rPVT from 30 days to 6 months after radiation exposure.

### Radiation Procedures

When rats were approximately 5-6 months old, they were exported to Brookhaven National Laboratory (BNL), seven days prior to the scheduled exposure day. All animals were exposed on the same day, and then returned to Johns Hopkins for follow-up testing 10-15 days post-irradiation. Rats were placed in individual plastic holders, and then acutely exposed to ^4^He ions (250 MeV/n) generated at the NASA Space Radiation Laboratory (NSRL) facility at BNL (sham: n=25, 5 cGy: n=29, and 25 cGy: n=29). ^4^He ions of this energy have a range in water of 37.6 cm with an average linear energy transfer (LET) of 1.58 keV/μm. Target exposure levels were 5 and 25 cGy; actual delivered doses varied by no more than 0.1 cGy relative to each target dose. Dose rates were approximately 5 cGy/min. The control group was sham irradiated (i.e., shipped to BNL, placed in the plastic holder, but not brought into the beam line).

### Social Odor Recognition Memory Test (SORM)

The SORM test consisted of three phases: familiarization, habituation, and the recognition test, each separated by 24 h and was conducted in a manner identical to our previous reports^11,48–50^. All phase 2 and 3 trials were video recorded with a digital video camera (Sony HDR CX240/L Digital Camcorder) and scored offline with Ethovision software (v.17.0, Noldus, Leesburg, VA) by experimenters blinded to the bead and rat irradiation conditions. Three SORM tests were completed following radiation exposure (1 month, 3 months, and 6 months; data for the 7-day SORM test from these rats was published previously)^48^. Different novel social odors from conspecifics housed in a different vivarium were used as odor donors. The phases below started seven days after the last cage cleaning, such that cages were seven days dirty prior to any beads being placed into the cages.

### Phase 1: Familiarization

Seven 2.5-cm round, unfinished wooden beads (CraftParts Direct, www.amazon.com) were introduced into each rat’s home cage to acquire the odor of individual subject rats and serve as familiar odors (beads F1, F2 and F3) for subsequent phases. Wooden beads were also placed into the cages of separate ‘‘odor donor’’ rats to provide novel 1 (N1) and novel 2 (N2) bead odors for subsequent test phases.

### Phase 2: Habituation

24 h after the familiarization phase and 1 h before testing, all wooden beads were removed and placed in individual plastic bags. For testing, three familiar odor beads and one novel 1 odor bead (N1) were introduced into each rat’s home cage. The order of the four beads was randomly altered for each rat. Beads were placed in a row in the front of each rat’s cage. Three 1-min trials separated by 1-min inter-trial intervals were completed. During each 1-min trial, rats were allowed to freely explore the beads. The first contact with any bead started each 1-min trial. All beads from this phase were discarded after use.

### Phase 3: Recognition Test

In a manner similar to the habituation phase, rats were presented with four beads 24 h following the habituation phase: two familiar odor beads, one novel odor 1 (N1) bead (the odor experienced during the habituation phase 24 h earlier) and one novel odor 2 (N2) bead (an unfamiliar, novel odor not experienced previously), with the order of the four beads randomly altered for each rat, for three 1-min trials with a 1-min interval between trials identical to the method described above for habituation. All rats readily approached the beads in both phases within a few seconds; however, a criterion of 2 min was adopted, such that if any rat did not begin exploring the beads within 2 min of their placement into the cage on any trial, the trial was concluded, and the beads were removed. This trial would be excluded from all calculations. No rat met this criterion at any time point throughout the study.

For each SORM test, time spent exploring [e.g., sniffing, whisking near (within 1 cm), manipulating beads with paws and/or mouth, licking beads] each wooden bead (familiar beads: F1-F3, N1 and N2) was measured in seconds for each 1-min trial for both the habituation and recognition test phases and summed separately for each trial. Percentage time exploring was calculated for each bead: time exploring bead/total time exploring all beads, e.g., N1/(N1 + F1 + F2 + F3). To compare the different groups on the recognition test at each time point, a discrimination index (DI) was also calculated using total exploration time summed separately for the N2, N1 and familiar beads as follows: DI = (N2 – N1)/(N2 + N1 + F1 + F2) * 100.

Discrimination index values greater than zero demonstrate significant preference for the N2 odor bead, whereas values less than zero show preference for the N1 odor bead. In addition, DI values equal to zero demonstrate that the N1 and N2 odor beads were explored to a similar degree.

#### Tissue collection

At 7, 30, 90-, and 180-days following radiation exposure, a subset of rats was euthanized to collect tissues for subsequent analysis (see Table 1). Ninety-minutes following the Recognition Test of the SORM, rats were euthanized via decapitation. Trunk blood was collected into EDTA-coated tubes and spun at 1500xg for 10 min at 4C, following which plasma was removed, snap-frozen, and stored at -80C until processing. Right hind limbs were removed, separated at the knee, cleaned of non-osseous tissue, and stored in 70% ethanol before transport to The College of New Jersey for testing.

**Table 1.** Group sizes for tissue collection

| Time Following<br>Radiation | <sup>4</sup>He Dose (cGy) |  |  |
| --- | --- | --- | --- |
|  | 0 | 5 | 25 |
| 7 days (1 week) | 5 | 5 | 5 |
| 30 days (1 month) | 5 | 5 | 5 |
| 90 days (3 months) | 5 | 8 | 8 |
| 180 days (6 months) | 10 | 11 | 11 |
| **Total number of rats** | **25** | **29** | **29** |

### Rat V-Plex Proinflammatory Cytokine Panel

Cytokine assays were completed using the Meso Scale Diagnostics (MSD) V-Plex Proinflammatory Panel 2 Rat Kit (Cat. No. K15059D, MSD, Rockville, MD), which allows for the simultaneous measurement of IFN-γ, IL-10, IL-13, IL-1β, IL-4, IL-5, IL-6, KC/GRO, TNF-α. Plasma aliquots were thawed and each sample was run in duplicate. 150µl of Blocker H was added to each well on the 96-well plate and incubated at room temperature for 1-hr with gentle shaking. The plate was then washed three times with PBS with Tween-20 (0.5%; PBS-T) wash buffer. 50µl of each sample (diluted in Diluent 42) or calibrator were added to each well, and again incubated at room temperature for 2-hrs with gentle shaking. The plate was then washed three times with PBS-T wash buffer. 25µl of detection antibody solution was added to each well. The plate was then incubated at room temperature for 2-hrs with gentle shaking. Finally, the plate was washed three times with PBS-T wash buffer. 150µl of MSD Read Buffer T was added to each well, and the plate was read immediately by the Sector Imager (MSD, Rockville, MD).

### Bone Mechanical Properties

Right femora were placed in saline solution to rehydrate 24-hours prior to testing. An Instron Universal Testing System (model 5967) was used to conduct three-point bend and femoral neck testing. During three-point bend, femora were tested in the anterior-posterior position until failure over a 10 mm span at a deflection rate of 0.05 mm/s. Femurs were supported below the femoral neck and above the condyles. A constant span length was chosen based on the most uniform area of the bone in which a point load, measured by a 500 N load cell, was applied in the center of the span. Force and deflection data was collected in order to quantify the mechanical properties of the femur (i.e. Linear Flexural Stiffness, Maximum Load, Energy to Failure).

After completion of three-point bend testing, the femoral neck portion of the bone was embedded in epoxy with the greater trochanter and femoral head placed upward. Bones were rehydrated in saline at least 24-hours prior to femoral neck testing. During femoral neck tests, the Instron was equipped with a 3.95 mm diameter cupped tip aligned to ensure that the cup was centered over the femoral head. The loading rate for the femoral neck was identical to the three-point bending analysis. The femoral neck was tested until failure using a 500 N load cell to measure force, deflection was also collected.

Force and deflection data from three-point bend and femoral neck testing was analyzed by a custom MATLAB code to quantify bone mechanical properties. Yield point was determined using the 90% secant method and was subsequently used to calculate additional mechanical values. The moving average of the linear region was utilized to determine linear flexural stiffness of the bone. Maximum and failure load were calculated using the largest y and x value, respectively. Energy to yield, maximum, and failure load were obtained by using a trapezoidal numerical integration from specific points on the curve (i.e., maximum load and failure load points).

### Data analysis

For the rPVT, the daily session averages for rPVT performance measure were used for subsequent analyses. A fully-saturated ANCOVA-type linear mixed effects (LME) model was fit to the rPVT data set to evaluate longitudinal performances for each performance measure (see Table 2). The fixed effects of the LME model were radiation dose, time, and the radiation dose x time interaction. The subject-specific intercept and slope served as the random effects of the LME. The sham control group was set as the reference level (Dose = 0). For each rPVT response variable, significant effects of radiation exposure were determined using two contrast comparisons (sham vs. 5 cGy and sham vs 25 cGy). Effects were considered significant if p-values were less than 0.05. The overall performance score (OPS; described above) was used for correlation analysis between rPVT performance and various tissue targets.

**Table 2.** LME Model Summary Table for rPVT Measures Following Helium Ion Exposure.

| rPVT<br>Response<br>Measure | Slope<br>contrast | Estimate | Std. Error | T-statistic | P-value |
| --- | --- | --- | --- | --- | --- |
| **Percent**<br>**correct** | 5-0 | -0.0035562 | 0.0045021 | -0.7898922 | 0.6404162 |
|  | 25-0 | -0.0218101 | 0.0045113 | -4.8345156 | **0.0000027\*** |
| **False**<br>**alarms** | 5-0 | 0.0041219 | 0.0038163 | 1.0800834 | 0.4467985 |
|  | 25-0 | 0.0117204 | 0.0038241 | 3.0648542 | **0.0042123\*** |
| **Premature**<br>**Responses** | 5-0 | -0.0013466 | 0.0008012 | -1.6807869 | 0.1619504 |
|  | 25-0 | -0.0003384 | 0.0008028 | -0.4215769 | 0.8769551 |
| **Lapses** | 5-0 | 0.0033978 | 0.0043795 | 0.7758554 | 0.6501061 |
|  | 25-0 | 0.0214964 | 0.0043884 | 4.8984412 | **0.0000019\*** |
| **Q10 RT** | 5-0 | 0.121608 | 0.0229732 | 5.2934739 | **0.0000002\*** |
|  | 25-0 | 0.1261701 | 0.0230205 | 5.4807802 | **0.0000001\*** |
| **Median RT** | 5-0 | 0.2519299 | 0.0430507 | 5.8519401 | **0.0000000\*** |
|  | 25-0 | 0.1297934 | 0.0431393 | 3.0087067 | **0.0050633\*** |
| **Q90 RT** | 5-0 | 0.1987873 | 0.1173013 | 1.6946724 | 0.1575752 |
|  | 25-0 | 0.5422591 | 0.1175423 | 4.6133097 | **0.0000079\*** |
| **Mean RT** | 5-0 | 0.2031358 | 0.0442344 | 4.5922588 | **0.0000087\*** |
|  | 25-0 | 0.2341631 | 0.0443254 | 5.2828198 | **0.0000003\*** |

**Note:** \*significant difference in slope between sham and exposed group.

To assess SORM performance within each group for 30, 90, and 180 day time points (the 7-day timepoint was published previously)^48^, separate paired t-tests were used to evaluate 1) exploration of the N1 odor compared to the familiar odors (F1 – F3) on Trial 1 of Habituation, 2) exploration of the N1 odor on Trial 3 of Habituation compared to Trial 1, and 3) exploration on the N2 odor compared to the N1 odor on the recognition test. To evaluate the effects of 5 and 25 cGy ^4^He exposure on SORM DIs at the different time points, a mixed-effects model (Radiation Dose x Time Following Radiation) with the Greenhouse-Geisser correction was used, followed by Dunnett’s multiple comparisons to assess specific group differences.

For IFN- γ, IL-13, IL-10, IL-4, IL-5, IL-6, KC/GRO, TNF-α, and undercarboxylated osteocalcin (ucOC), separate two-way ANOVAs were used to assess the main-effects of Radiation Dose and Time Following Radiation, and the Radiation Dose x Time Following Radiation interaction. Significant effects were followed by Tukey’s multiple comparisons test. For IL-1beta, an independent-samples t-test was used to compare sham and 5 cGy at the 3-month timepoint, since too few 25 cGy samples had detectable levels of this cytokine. For the 6-month timepoint for IL-1beta, a one-way ANOVA followed by Tukey’s multiple comparisons test was used to compare levels of this cytokine between the three groups. Spearman’s rank correlation was used to measure the correlation between behavioral parameters (DI, rPVT OPS) and cytokines or ucOC. All analysis were completed with SPSS Statistics (v28) or GraphPad Prism (v10.1.0).

Analysis of bone mechanical properties was performed using Kruskal-Wallis and Dunn pairwise comparison tests were performed in R-Studio (4.2.2), with an alpha=0.05. Principal Component Analysis was also performed in R-Studio (4.2.2) to compare the trends in bone properties, cytokine measurements, and neurobehavioral performance.

## Results

### Baseline (pre-radiation) sustained attention is similar between groups

To determine if pre-radiation rPVT performance was equivalent between the three groups (0 cGy, 5 cGy, and 25 cGy), mean rPVT behavioral parameters for the two weeks prior to radiation exposure were analyzed with one-way ANOVAs. No significant differences between the groups were found for any measure prior to radiation exposure (all *p’*s ≥ 0.3363; Figure 1, left panels).

**Figure 1.**
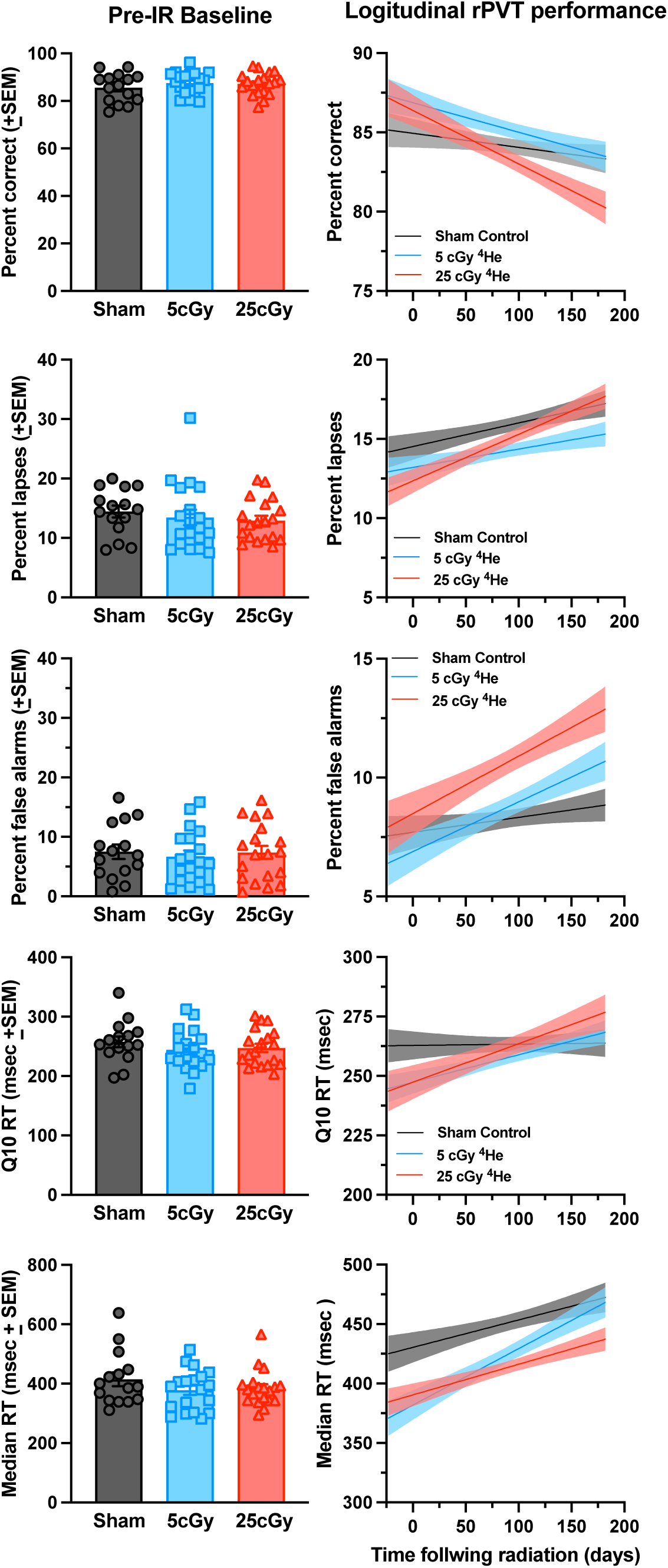
rPVT performance parameters at the pre-irradiation baseline (left panels) and throughout the post-exposure testing period (right panels). A) Mean percent correct responding. B) Mean percent lapses in attention. C) Mean percent false alarms. D) Mean Q10 reaction times. E) Median reaction times. See Table 2 for what measures significantly different between the groups.

### 4He exposure dose-dependently impairs sustained attention

The rPVT is a rodent test of sustained attention, and significant changes on various response parameters suggest that radiation exposure could impair attention and response times in a dose-dependent manner. Specifically, 25 cGy ^4^He exposure resulted in a significant decrease in correct responding across the post-radiation period compared to the sham control group (*p* < 0.0001; Table 2), but 5 cGy ^4^He exposure did not affect correct responding compared to the sham control group (*p* = 0.64). False alarms (premature responses during the 3-10 sec window when rats wait for the onset of the stimulus light) and lapses in attention were significantly increased in the 25 cGy group compared to sham controls (*p*’s ≤ 0.004), but no effect of 5 cGy ^4^He exposure was found on these response parameters. However, both doses of ^4^He ions significantly increased several response latency parameters, including Q10 response times (i.e., 10^th^ percentile reaction times or the fastest 10% of reaction times), median response times, and mean response times. In general, exposure to ^4^He ions, regardless of the dose, slowed most response times on the rPVT (see Table 2).

### 4He exposure impairs social odor recognition memory

The SORM test was used to assess the effects of radiation on recognition memory based on social odors (i.e., same-sex conspecifics). During Habituation, all groups regardless of radiation exposure and time following radiation, displayed a significant preference for the Novel 1 odor compared to the Familiar odors (see Table 3). Further, each group decreased exploration of the Novel 1 odor by the third trial, demonstrating habituation to this odor. For the Recognition Test, sham control rats displayed significantly greater exploration of the Novel 2 odor compared to the Novel 1 odor for each time point, which supports intact social recognition memory over time in these animals. However, both doses of ^4^He radiation resulted in similar levels of exploration of the Novel 2 and Novel 1 odors at each time point tested, which suggests that ^4^He radiation significantly impaired social recognition memory starting at 30 days following exposure with no recovery of social odor recognition memory up to 6 months following exposure.

**Table 3.** Exploration in the SORM at each timepoint following exposure.

| **Habituation** |  |  |  |  |  |  |  |
| --- | --- | --- | --- | --- | --- | --- | --- |
| **30 Days** | **N1 Trial 1** | **Familiar Trial 1** | **t value** | **p value** | **N1 Trial 3** | **t value** | **p value** |
| Sham | 12.98 ±1.89 | 3.68 ±0.59 | 4.991 | <0.001 | 4.18 ±1.42 | 4.52 | <0.001 |
| 5 cGy | 12.94 ±2.28 | 4.25 ±0.62 | 3.558 | 0.002 | 4.94 ±1.63 | 3.739 | 0.001 |
| 25 cGy | 12.28 ±1.77 | 4.59 ±0.49 | 4.391 | <0.001 | 5.73 ±1.56 | 3.178 | 0.004 |
| **90 Days** | **N1 Trial 1** | **Familiar Trial 1** | **t value** | **p value** | **N1 Trial 3** | **t value** | **p value** |
| Sham | 13.14 ±1.87 | 4.83 ±0.69 | 4.918 | <0.001 | 5.96 ±2.09 | 2.727 | 0.016 |
| 5 cGy | 13.12 ±2.58 | 4.38 ±0.59 | 3.818 | 0.002 | 4.21 ±1.36 | 4.182 | <0.001 |
| 25 cGy | 22.56 ±3.69 | 4.11 ±0.51 | 4.949 | <0.001 | 13.16 ±3.88 | 2.368 | 0.031 |
| **180 Days** | **N1 Trial 1** | **Familiar Trial 1** | **t value** | **p value** | **N1 Trial 3** | **t value** | **p value** |
| Sham | 15.26 ±4.45 | 4.10 ±0.73 | 2.275 | 0.049 | 8.28 ±4.18 | 1.600 | 0.144 |
| 5 cGy | 21.62 ±3.85 | 3.25 ±0.80 | 4.564 | 0.001 | 12.43 ±5.35 | 2.552 | 0.029 |
| 25 cGy | 17.09 ±3.71 | 4.28 ±0.95 | 3.443 | 0.007 | 9.52 ±4.72 | 2.104 | 0.062 |
| **Recognition** |  |  |  |  |  |  |  |
| **30 Days** | **N1 Trial 1** | **N2 Trial 1** | **t value** | **p value** |  |  |  |
| Sham | 21.90 ±4.02 | 46.19 ±4.58 | 3.238 | 0.005 |  |  |  |
| 5 cGy | 37.04 ±6.82 | 44.94 ±6.54 | 0.68 | 0.504 |  |  |  |
| 25 cGy | 27.00 ±5.95 | 44.85 ±5.98 | 1.575 | 0.13 |  |  |  |
| **90 Days** |  |  |  |  |  |  |  |
| Sham | 5.77 ±0.89 | 16.96 ±3.36 | 3.549 | 0.003 |  |  |  |
| 5 cGy | 11.74 ±2.22 | 14.94 ±3.33 | 0.705 | 0.49 |  |  |  |
| 25 cGy | 9.52 ±1.66 | 15.71 ±3.92 | 1.283 | 0.216 |  |  |  |
| **180 Days** |  |  |  |  |  |  |  |
| Sham | 4.90 ±1.09 | 22.50 ±5.32 | 3.042 | 0.014 |  |  |  |
| 5 cGy | 14.86 ±4.28 | 11.79 ±1.47 | 0.207 | 0.841 |  |  |  |
| 25 cGy | 13.53 ±4.61 | 14.80 ±5.47 | 0.083 | 0.936 |  |  |  |

A mixed-effects model (Radiation Dose x Time Following Radiation) was used to compare the DI values for all time points for the irradiated and sham control groups. The main-effect of Radiation Dose was significant [F(2,135) = 5.641, *p* = 0.0044], but the main-effect of Time Following Radiation (*p* = 0.8207) and the Radiation Dose x Time Following Radiation interaction (*p* = 0.9127) were not significant (Figure 2A). Post-hoc analyses identified that both irradiated groups had DI values that were significantly lower than that of the sham control group, but they did not differ from each other (all *p*’s ≤ 0.0306; Figure 2B). Thus, ^4^He exposure impaired social odor recognition memory in a dose- and time-independent manner, irrespective of when it was measured during the post-exposure period (from 30 to 180 days).

**Figure 2.**
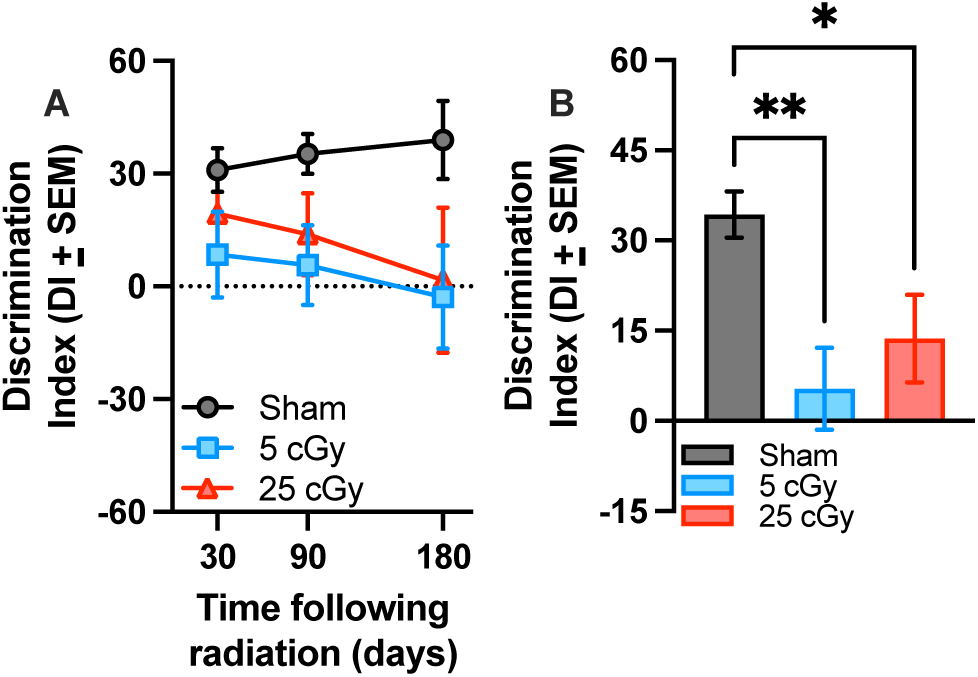
^4^He (250 MeV/n) significantly impairs social odor recognition memory compared to unexposed sham control rats. A) Mean SORM discrimination index (DI ± SEM) for sham exposed (gray), 5 cGy (blue), and 25 cGy (red) ^4^He exposure groups. B) Mean SORM discrimination index (DI + SEM) for all groups collapsed across time representing the significant main-effect of Radiation Dose. Both doses of ^4^He resulted in significant decreases in social odor recognition memory compared to sham exposed rats. \**p* < 0.05, \*\**p* < 0.01.

### Plasma TNF-α, IL-1β, and ucOC levels correlate with rPVT performance

A significant main-effect of Radiation Dose was found for TNF-α levels [F(2,46) = 3.999, *p* = 0.0250], with the 25 cGy exposure group displaying lower plasma levels of this cytokine compared to the 5 cGy exposure group (*p* = 0.0293; Figure 3 top); however, this group did not differ significantly from the sham exposure group (*p* = 0.1075), nor did the 5 cGy group (*p* = 0.8689). Spearman correlation was used to evaluate the relationship between rPVT performance using the OPS and TNF-α levels at the 3- and 6-month time points. A near-significant negative correlation [*r*(9) = -.58, *p* = 0.0656] was found in the sham control rats between rPVT OPS and TNF-α levels. An opposite, near-significant positive correlation [*r*(15) =.439, *p* = 0.07] was found between the rPVT OPS and plasma TNF-α levels in the 5 cGy-exposed rats, with greater rPVT OPS (better performance) associated with higher levels of plasma TNF-α. No relationship was found between rPVT OPS and TNF-α levels in the 25 cGy group. At the 6-month timepoint, the one-way ANOVA evaluating plasma IL-1β levels was significant [F(2, 10) = 7.337, *p* = 0.0109], and the 25 cGy exposure group displayed significantly lower plasma levels of this cytokine compared to both sham controls and the 5 cGy exposure group (all *p’*s ≤ 0.0330; Figure 3 bottom). There were no differences between sham controls and the 5 cGy exposure group (*p* = 0.6783). A significant positive correlation was found for rPVT OPS and plasma IL-1β levels at this timepoint [*r*(11) = .584, *p* = .040] with greater rPVT OPS (better performance) associated with higher plasma IL-1β levels. No significant effects were found for the remaining cytokines (i.e., IL-1β at the 3-month time point, IFN-gamma, IL-13, IL-10, IL-14, IL-5, IL-6, and KC/GRO, all *p’*s ≥ 0.0539). A significant main-effect of Timepoint was found for plasma ucOC levels [F(3, 77) = 9.135, *p* <0.001], but not Radiation Dose or their interaction (all *p’*s ≥ 0.302; Figure 4). Plasma ucOC levels were significantly lower at 7-days compared to levels measured at 30-days (*p* = 0.025) and 90-days (*p* <0.001), but were comparable to levels measured at 180-days (*p* = 0.136). Plasma levels at 180-days were also significantly lower than levels measured at 90-days (*p* = 0.003). A near-significant positive correlation was found for rPVT OPS and plasma ucOC levels [*r*(51) = .2633, *p* = 0.056], with greater rPVT OPS associated with higher plasma ucOC levels.

**Figure 3.**
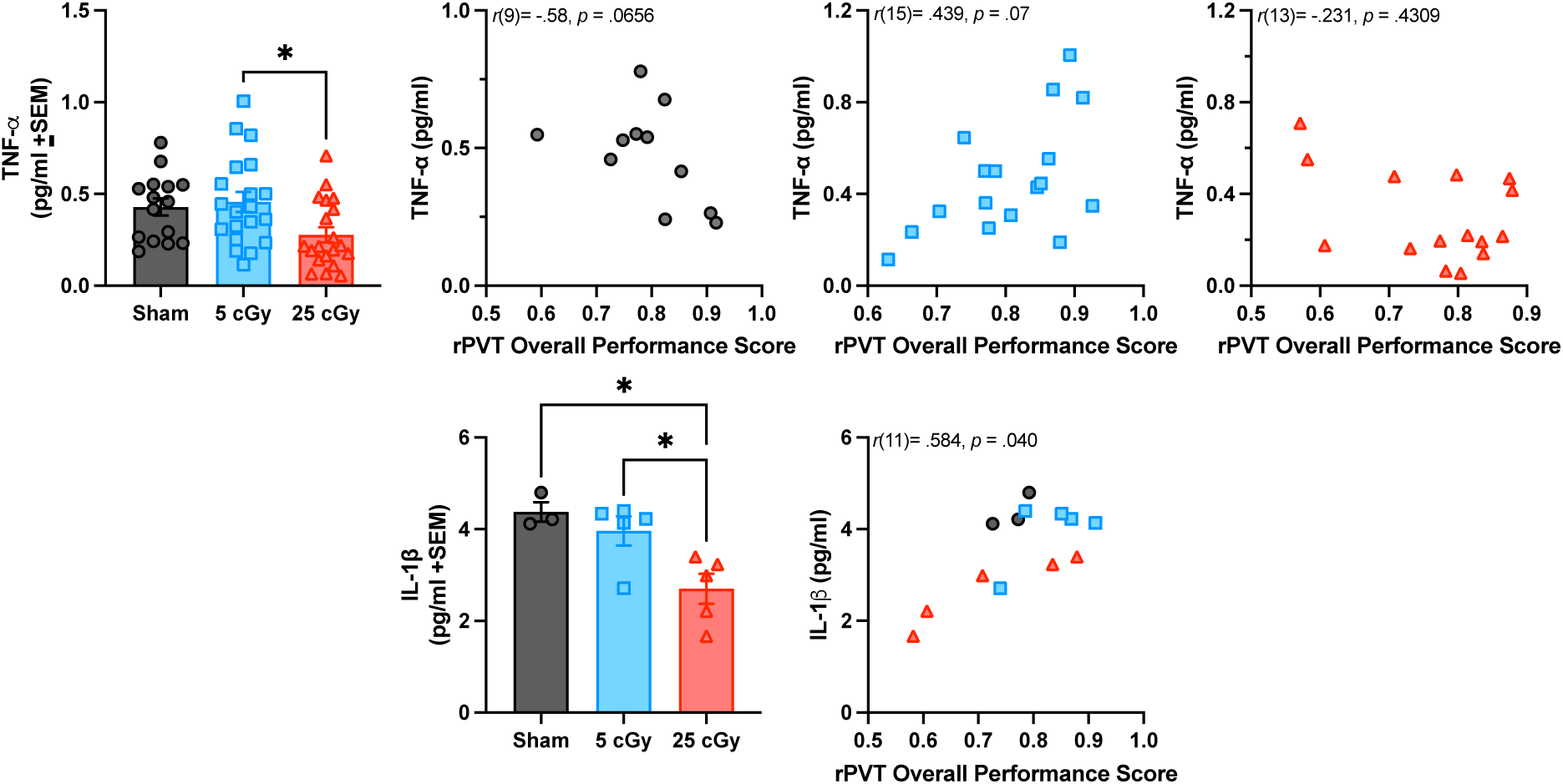
Mean plasma IL-1 and TNF-α levels in irradiated and sham control rats measured at 3 and/or 6 months following ^4^He exposure. Top: TNF-α was significantly lower in the 25 cGy-exposed group compared to the 5 cGy-exposed group. For sham rats, TNF-α levels were negatively correlated with rPVT OPS. For 5 cGy-exposed rats, a positive correlation was found for TNF-α levels and rPVT OPS score. Bottom: At 6-months following exposure, a significant decrease in plasma IL-1b levels was found only in the 25 cGy group. A significant correlation was found between plasma IL-1b levels and rPVT OPS, with greater OPS associated with higher levels of this cytokine. \**p* < 0.05.

**Figure 4.**
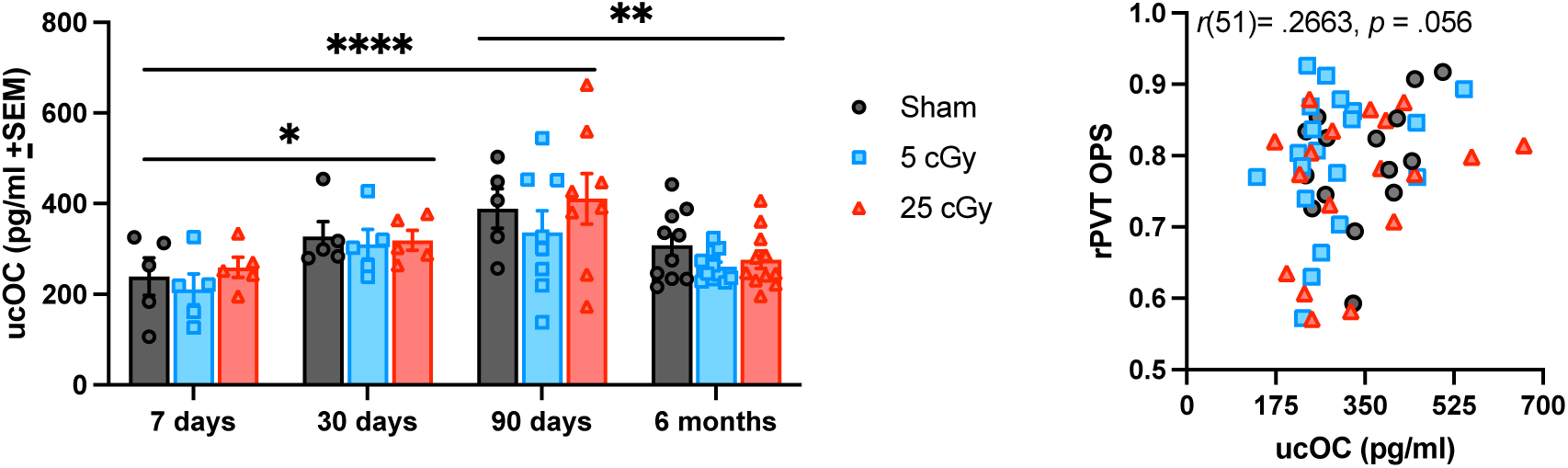
Left: Mean plasma undercarboxylated osteocalcin (ucOC) levels in irradiated and sham control rats measured at different timepoints following exposure. Significant effects of Time were found, but no effects of radiation. \**p*<0.05; \*\**p*<.01; \*\*\*\**p*<0.0001. Right: Near-significant positive correlation between ucOC level and rPVT OPS for rats in the 90 day and 6-month groups. Better rPVT performance was correlated with higher plasma levels of ucOC.

### Bone mechanical properties differ by timepoint, but not radiation exposure

For both 3-point bend and femoral neck testing there was a main-effect of Time Following Radiation but not Radiation Dose (see Table 4). For 3-point bend testing, effects of Time Following Radiation occurred primarily in measures at Yield with no effect in measures at Max Load or Failure (all *p’*s ≥ 0.093, Table 4). Stiffness generally increased over time and was significantly higher at 180-days compared to 7-days (*p* = 0.008, Fig 5A) with near-significance by 90-days (*p* = 0.033). Between 90- and 180-days there was a decrease in Displacement at Yield (*p* = 0.023, Fig 5B) and Energy to Yield (*p* = 0.014, Fig 5C), but an increase in Post-Yield Displacement (*p* = 0.003, Fig 5D).

**Figure 5.**
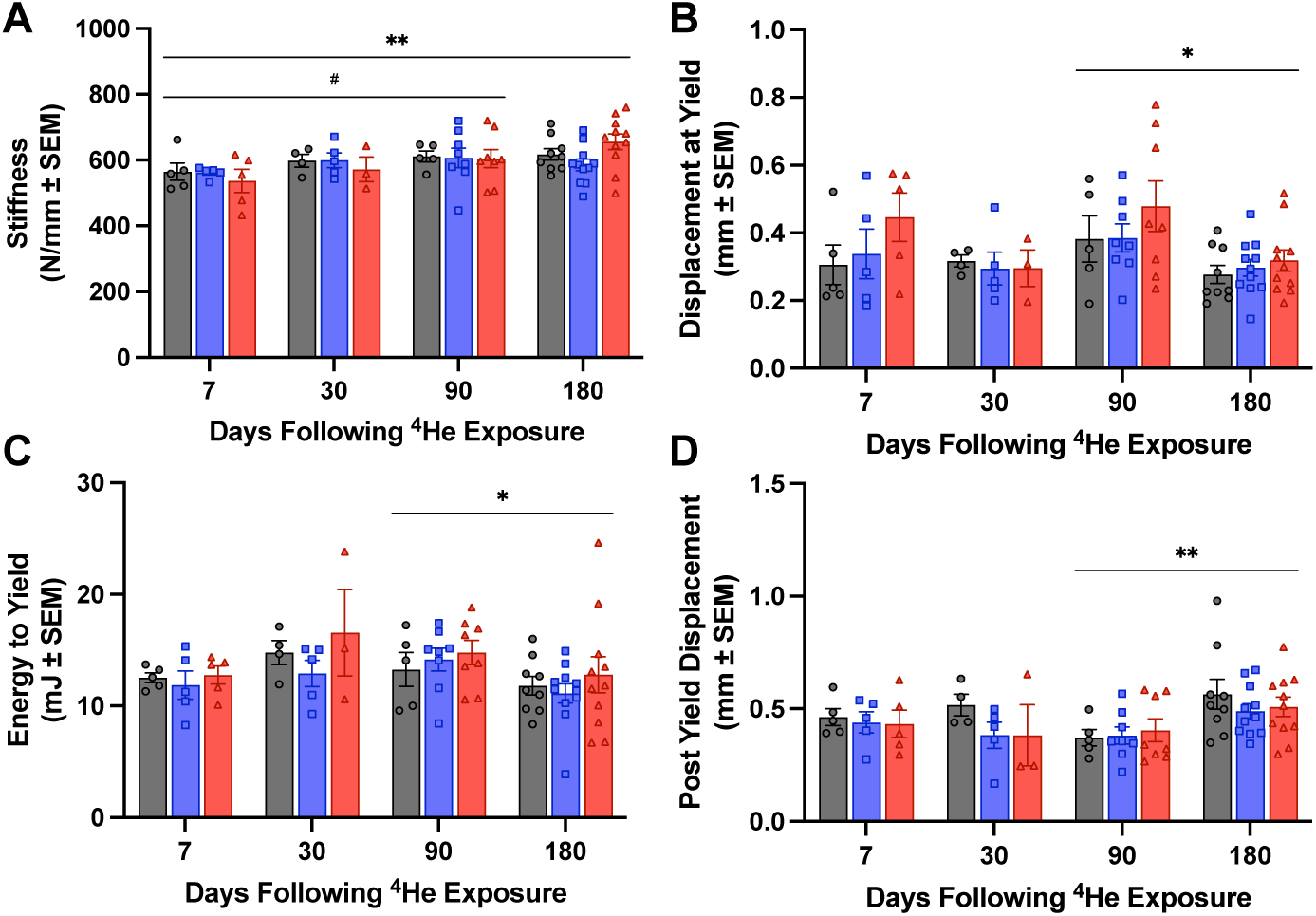
Three-point bend measures of bone strength over time, including: A) Stiffness, B) Displacement at Yield, C) Energy to Yield, and D) Post-Yield Displacement. \**p* < 0.025; \*\**p*<0.01; #*p*=0.033. Gray = Sham control; Blue = 5 cGy; Red = 25 cGy.

**Table 4.** Summary of *p*-values for main-effect and pairwise comparisons as measured for bone biomechanical properties.

|  | Main-Effect |  | Pairwise Comparison for Time |  |  |  |  |  |
| --- | --- | --- | --- | --- | --- | --- | --- | --- |
|  | Dose | Time | 7D vs 30D | 7D vs 90D | 7D vs 180D | 30D vs 90D | 30D vs 180D | 90D vs 180D |
| **3-point Bend Test** |  |  |  |  |  |  |  |  |
| Stiffness (N/mm) | 0.5177 | **0.019\*** | 0.384 | 0.033 | **0.008\*** | 1.000 | 0.862 | 1.000 |
| Displacement at Yield (mm) | 0.2594 | **0.049\*** | 1.000 | 0.609 | 0.904 | 0.152 | 1.000 | **0.023\*** |
| Yield Load (N) | 0.6824 | **0.038\*** | 0.147 | 0.076 | 1.000 | 1.000 | 0.223 | 0.102 |
| Energy to Yield (mJ) | 0.6936 | **0.013\*** | 0.356 | 0.204 | 1.000 | 1.000 | 0.061 | **0.014\*** |
| Post-Yield Displacement (mm) | 0.5222 | **0.012\*** | 1.000 | 0.440 | 0.508 | 0.792 | 0.381 | **0.003\*** |
| Displacement at Max Load (mm) | 0.648 | 0.3071 | - | - | - | - | - | - |
| Max Load (N) | 0.6792 | 0.2939 | - | - | - | - | - | - |
| Energy to Max Load (mJ) | 0.822 | 0.4124 | - | - | - | - | - | - |
| Displacement at Failure (mm) | 0.397 | 0.6305 | - | - | - | - | - | - |
| Failure Load (N) | 0.8873 | 0.09326 | - | - | - | - | - | - |
| Energy to Failure (mJ) | 0.3678 | 0.1867 | - | - | - | - | - | - |
| **Femoral Neck Test** |  |  |  |  |  |  |  |  |
| Stiffness (N/mm) | 0.7677 | **< 0.001\*** | 0.496 | 0.071 | 0.057 | **0.001\*** | **< 0.001\*** | 1.000 |
| Displacement at Max Load (mm) | 0.8301 | **< 0.001\*** | 1.000 | 0.099 | **< 0.001\*** | 0.176 | **0.001\*** | 0.297 |
| Max Load (N) | 0.636 | 0.3822 | - | - | - | - | - | - |
| Energy to Max Load (mJ) | 0.8404 | **< 0.001\*** | 1.000 | 0.296 | **< 0.001\*** | 0.050 | **< 0.001\*** | 0.144 |
| Displacement at Failure (mm) | 0.8522 | **< 0.001\*** | 1.000 | 0.085 | **< 0.001\*** | 0.105 | **< 0.001\*** | 0.380 |
| Failure Load (N) | 0.4834 | 0.3964 | - | - | - | - | - | - |
| Energy to Failure (mJ) | 0.8104 | **< 0.001\*** | 1.000 | 0.195 | **< 0.001\*** | 0.029 | **< 0.001\*** | 0.217 |

**Notes:** \*indicates significances based on either Kruskal-Wallis ( $p < 0.05$ ) for main-effect analysis or Dunn ( $p < 0.025$ ) for pairwise comparisons.
Factor Abbreviations: Dose = Radiation Dose; Time = Time After Radiation

At the femoral neck, a main-effect of Time Following Radiation was found for Femoral Neck Stiffness, Displacement at Max Load, Energy to Max Load, Displacement at Failure, and Energy to Failure (all *p’*s < 0.001, Table 4). Pairwise analysis showed Stiffness at the femoral neck increased sharply between 30- and 90-days (*p* = 0.001) and remained elevated at 180-days (*p* < 0.001) post exposure (Fig 6). In contrast, there was a significant decline between early (7- and 30-Day) and late (180-Day) timepoints in measures of Displacement at Max Load (*p* ≤ 0.001, Fig 7A), Energy to Max Load (*p* < 0.001, Fig 7B), Displacement at Failure (*p* < 0.001, Fig 7C), and Energy to Failure (*p* < 0.001, Fig 7D) at the femoral neck.

**Figure 6.**
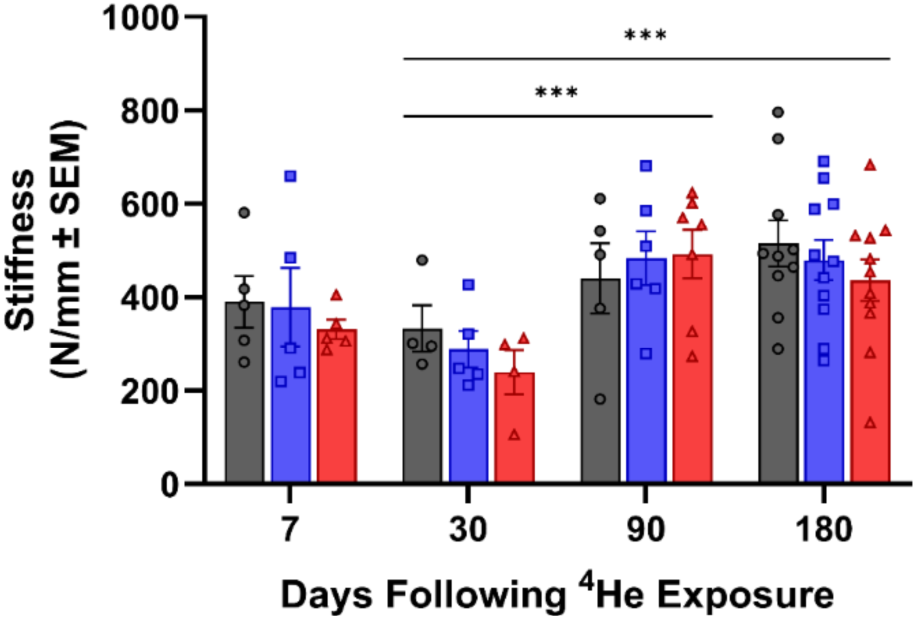
Bone stiffness varies over Time, but not Radiation Dose, at the femoral neck. ***p<0.001. Gray = Sham control; Blue = 5 cGy; Red = 25 cGy.

**Figure 7.**
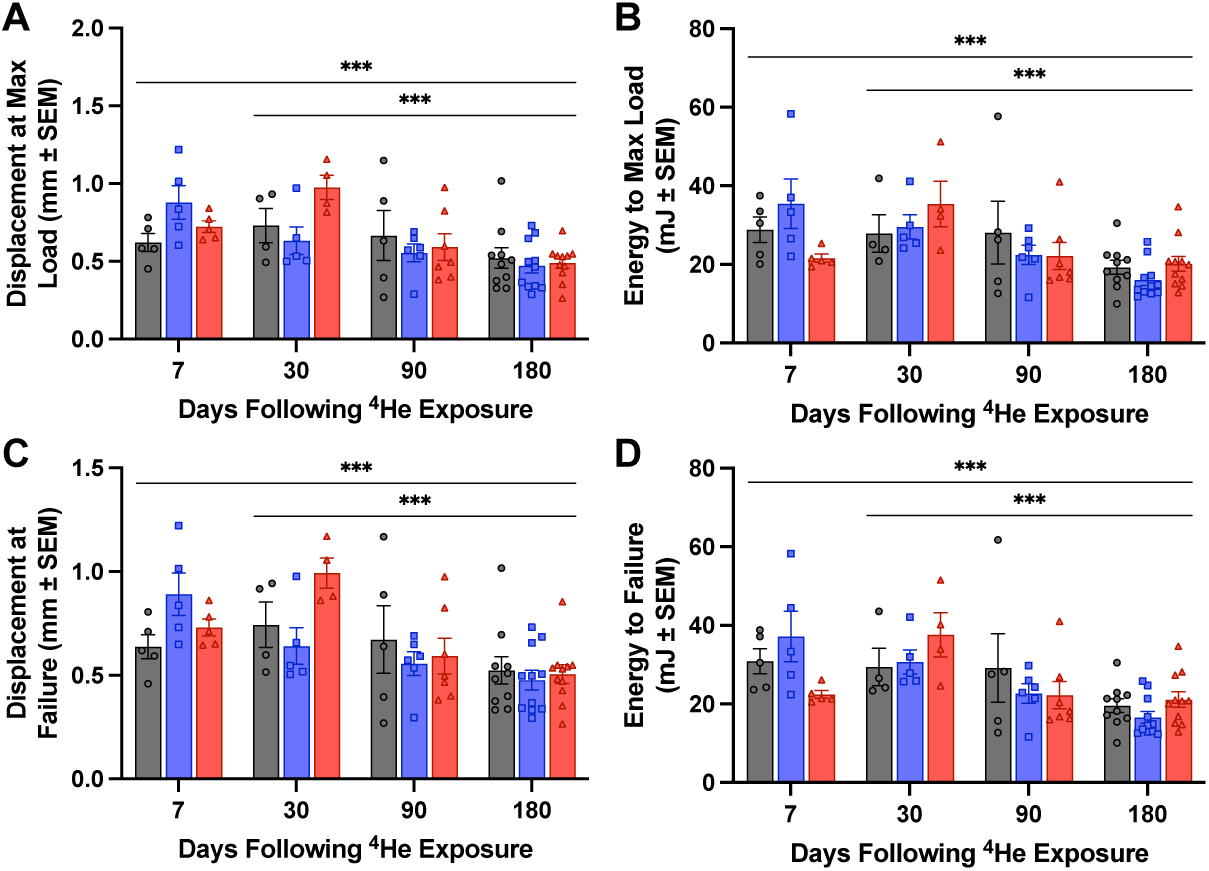
Additional measures of bone strength measured over time at the femoral neck, including: A) Displacement at Max Load, B) Energy to Max Load, C) Displacement at Failure, and D) Energy to Failure. *p<0.001. Gray = Sham control; Blue = 5 cGy; Red = 25 cGy.

### Relationships between CNS & Bone Property Changes

A combined dataset was created using all of the experimental data from the current study and our previously collected material property data and 7-day SORM performance from these same animals^49,52^. A Principal Component Analysis (PCA) was conducted to explore the relationships among bone quality, microindentation, and biochemical markers. The first two principal components explained a total of 39.4% of the variance in the dataset, with Component 1 accounting for 23.7% and Component 2 accounting for 15.7%. Variables related to mechanical bone strength, including 3-Point Bending Max Load, 3-Point Bending Failure Load, 3-Point Bending Stiffness, and 3-Point Bending Energy to Max Load, showed strong positive loadings on Component 1, indicating that this axis reflects overall bone mechanical performance. Similarly, microindentation measures (Relaxed Elastic Modulus, REM; Instantaneous Elastic Modulus, IEM) loaded positively on Component 1, suggesting alignment between microstructural integrity and mechanical performance. In contrast, variables such as 3-Point Bending Displacement at Yield, 3-point Bending Displacement to Failure, KC_GRO, and ucOC contributed more to Component 2, which may capture variation associated with biochemical or growth-related factors. Variables such as the microindentation measures and 3-point bending Max Load demonstrated high cos² values, implying that these variables are well represented in the reduced two-dimensional space. These results suggest that bone mechanical properties and microindentation metrics contribute significantly to the underlying structure of the data, with additional variation captured by biochemical markers along a secondary axis A Spearman’s correlation was also run on this larger data set to evaluate relationships between bone, behavioral, and cytokine measures. When taken as one dataset without respect to Radiation Dose or Time Following Radiation, biomechanical (3-point bend and femoral neck) and material property measures correlated with behavioral and cytokine data across several measures, although the relationships appeared to be in opposite directions (Fig 8). SORM DI was negatively correlated with 3-point bend Yield Load (r[77] = -.285, *p* = 0.012) and Energy to Yield (r[77] = -.301, *p* = 0.008). In contrast, there was a near positive correlation between SORM DI and IEM (r[79] = .220, *p* = 0.052) and REM (r[79] = .207, *p* = 0.067), both measures of bone material. Among 3-point bend data, Displacement at Yield correlated with several cytokines including IL-1β (r[28] = -.447, *p* = 0.017), IL-10 (r[63] = -.412, *p* = 0.001), and KC_GRO (r[76] = .274, *p* = 0.017) and there was an additional weak correlation with TNF-α (r[63] = -.243, *p* = 0.055). Displacement at Max Load correlated with IL-1β (r[28] = -.406, *p* = 0.032), IL-10 (r[63] = -.429, *p* < 0.001), and, weakly, with KC_Gro (r[76] = .221, *p* = 0.055). Displacement at Failure negatively correlated with IL-10 (r[63] = -.261, p = 0.039). IL-6 correlated negatively with Energy to Failure at the femoral neck (r[56] = -.282, *p* = 0.035) and positively with IEM (r[59] = .268, *p* = 0.040) and REM (r[59] = .301, *p* = 0.021). There were also near correlations between IL-6 and Max Load (r[56] = -.260, *p* = 0.053), IL-6 and Energy to Max Load (r[56] = -.261, *p* = 0.052), and TNF-α and REM (r[64] = .245, *p* = 0.051). There were no correlations between ucOC and any bone parameters (all *p’*s ≥ 0.083).

**Figure 8.**
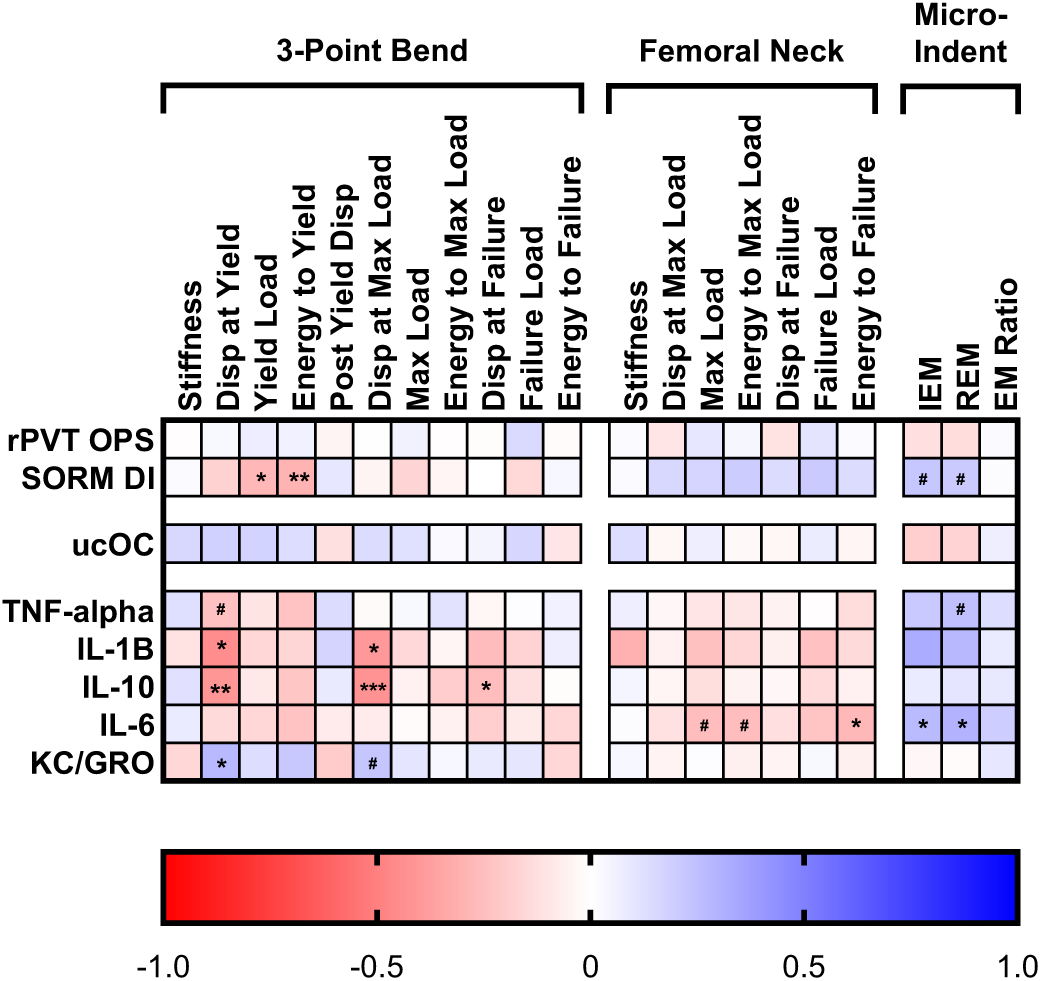
Heatmap of r values from Spearman’s correlation between the combined bone dataset and experimental data from behavioral testing and cytokine panels. *p<0.05, **p<0.01, ***p<0.001, #0.051≤p≤0.067.

Further analysis with linear regression revealed that some of the correlations vary with Radiation Dose. For Relaxed Elastic Modulus, the positive correlation is preserved in rats in the control group (r = .563, *p* = 0.004), but disappears in the 5 cGy (r = .336, *p* = 0.087) and 25 cGy (r = .067, *p* = 0.734) animals (Fig 9, Top). Interestingly, the SORM DI-Yield Load relationship behaves in the exact opposite manner with the correlation only occurring in the 25 cGy dose group (r = -.434, *p* = 0.236), but not in the 5 cGy (r = -.093, *p* = 0.674) or control (r = -.130, *p* = 0.517) animals (Fig 9, Bottom).

**Figure 9.**
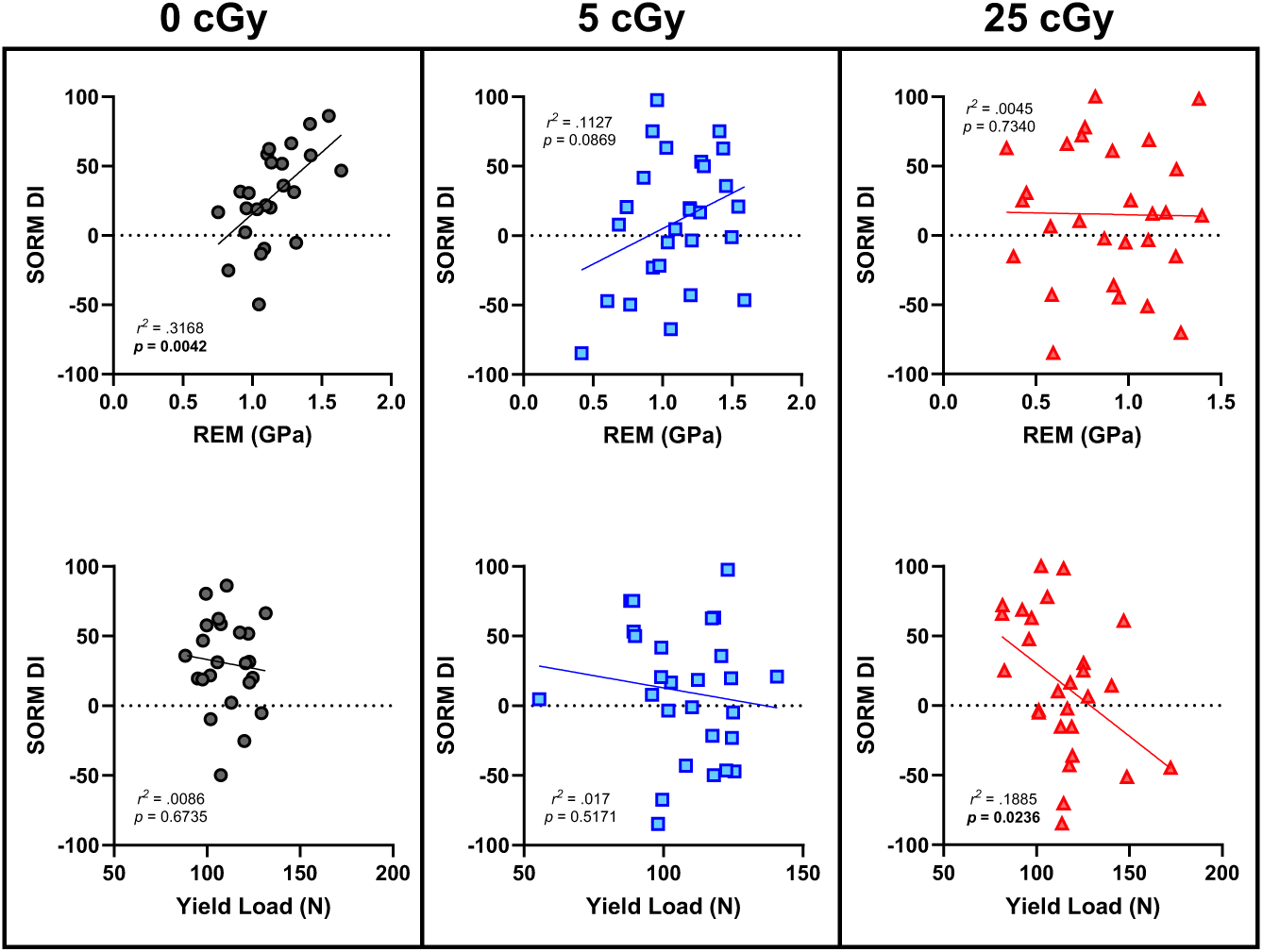
Linear regression of SORM DI and (Top) Relaxed Elastic Modulus (REM) and (Bottom) Yield Load.

## Discussion

We demonstrated that exposure to ^4^He ions, a major component of GCR, can significantly impair sustained attention and recognition memory when measured from 30- to 180 days following exposure, without any evidence of recovery of these functions. These low exposure doses of ^4^He, however, did not significantly alter bone mechanical properties measured in the same animals. Significant changes in skeletal integrity were noted as a function of time following exposure, which is likely an effect of age on bone mechanical properties, but may be exacerbated by the higher dose (25 cGy) radiation. We also found that circulating cytokines and ucOC levels were significantly correlated with behavioral performances, which suggests potential for these blood-based targets to serve as diagnostic biomarkers.

Radiation-induced deficits in sustained attention like those reported here would be concerning for astronauts for several reasons. First, sustained attention is a basic cognitive process and the ability to sustain attention to a task underlies more advanced skills associated with executive function; thus, deficits in sustained attention would likely permeate cognitive functioning in general and be apparent as greater distractibility, more errors, slower response times, and greater impulsivity across multiple tasks. For example, in a cohort of astronaut-like subjects, greater sustained attention, decreased response speed, and lower impulsivity were associated with better performance on an operationally-relevant 6-degrees-of-freedom docking task^51^, which suggests that high levels of sustained attention (measured as better performance on the PVT) are needed for superior performance of operationally-relevant tasks that utilize multiple cognitive domains. Second, the deficits found in the 25 cGy group appear to be similar to those found in rats performing this version of the rPVT, and others, following sleep fragmentation/disruption^46,52–54^. Specifically, sleep disruption in both rodents and humans increases attention lapses on the PVT, and in humans, is associated with increased accidents and performance errors^55,56^. Given that astronauts are known to experience disrupted sleep^57^, it is likely the effects of radiation compounded with sleep loss would increase these deficits, leading to an increase in errors, and potentially catastrophic accidents. Indeed, a recent study showed that increasing radiation exposure is associated with poorer PVT performance in astronauts on the International Space Station during a 6-month mission^44^. Third, we have previously shown that proton radiation exposure resulted in individual differences in rPVT deficits, such that some rats were seemingly unaffected by the exposure and displayed rPVT performance similar to sham controls^41^. However, in the present study, we have shown that ^4^He exposure affects subjects at the group level, such that on average, animals displayed decreasing performance across the post-radiation response period. Given that ^4^He ions are a large component of galactic cosmic rays (GCR), exposure to the GCR simulation would be expected to have similar (or more severe) effects as ^4^He ions on sustained attention, which is concerning for exploration-class missions.

Indeed, a recent study exploring the effects of acute or chronic exposure to the simplified GCR simulation (^1^H, ^4^He, ^16^O, ^28^Si, and ^56^Fe) showed that acquisition of an attention task was significantly delayed following both exposures in mice^58^. While this task did not measure lapses in attention like the rPVT, it provides support for GCR exposure leading to impaired sustained attention. However, it is possible that these group differences are unique to ^4^He exposure, which would still be disturbing for exploration-class missions, given the abundance of ^4^He in GCR. Finally, the lack of recovery during the 6-month post-radiation testing period and the fact that deficits appeared to worsen as the time post-exposure increased, suggests that deficits in sustained attention would be their most severe near the end of the mission, increasing risks associated with navigating the return to Earth.

Rats also displayed significant deficits in social recognition memory at all time points tested, which further supports lack of recovery of radiation-induced neurobehavioral deficits in a different cognitive domain. Exposure to a variety of space relevant ions, in addition to the GCR simulation, has supported deficits in recognition memory tasks, with most tasks using object identification (e.g., novel object recognition)^59^. However, time courses of these effects measured in the same subjects have not been presented, but the current results provide strong support for continued impairment in recognition memory in the same subject for at least 6 months following exposure. We previously reported an early (7 days following exposure), dose-dependent effect of ^4^He ions on social recognition memory^48^, but the 5 cGy group showed deficits equivalent to the 25 cGy at all following time points beginning at the 30-day time point presented here. Together, the behavioral data support long-term, progressive deficits in multiple cognitive domains following ^4^He ion exposure.

Identifying circulating markers of radiation effects that could be measured in astronauts is an important and on-going area of research. Several recent rodent studies have shown that various immune markers are associated with neurobehavioral performance following exposure in laboratory animals, such as T cells and monocytes^13,60,61^. We found significant decreases in circulating TNF-α and IL-1β in the 25 cGy exposed rats, suggesting that long-term neurobehavioral deficits are associated with an aberrant immune response. Interestingly, these cytokines were not altered in the 5 cGy exposed group, even though they had deficits in recognition memory and reaction time measures. Peripheral immune markers could be more closely related to changes in sustained attention, which would support these limited deficits at this lower dose. Indeed, spaceflight alters immune responses in astronauts, and radiation-induced changes in immune system function would likely worsen these effects. There were no radiation-induced changes in peripheral ucOC levels, only differences based on timepoint, which likely reflects age-related changes in blood levels of this hormone^62^.

These low exposure doses of ^4^He did not significantly alter the overall bone biomechanical response in these animals, however, specific individual parameters of bone strength were altered (e.g. Max Load, Displacement at Yield). Significant changes in skeletal integrity were noted as a function of time following exposure, which is likely an effect of age on bone mechanical properties but may be exacerbated by the higher dose (25 cGy) radiation. At higher doses (>50 cGy), skeletal deterioration in response to radiation is most likely due to a rapid increase in osteoclastic resorption^4,63–65^ and a corresponding loss of trabecular bone. The femoral midshaft, where 3-point bend testing is conducted, has almost no trabecular bone present, and bone’s strength characteristics at this location are primarily dependent on cortical properties. In contrast, the femoral neck contains both a cortical shell and an intricate network of trabecular bone to help distribute loads generated through normal activity. However, there are significant limitations on this test in terms of boundary conditions.

Previously we have reported a significant decline in bone material properties (Instantaneous & Relaxed Elastic Moduli; IEM, REM) in animals in this study exposed to 25 cGy vs controls at the 90-day timepoint^66^. In the current study there were also significant relationships between CNS and bone when looking at specific time points (90-Days after exposure). A significant correlation was found across all doses between SORM DI and REM (*p* = 0.042, Fig 10) and near-significant correlation between SORM DI and IEM (*p* = 0.053). Interestingly, this correlation appears to be dose dependent. In the non-irradiated control group, SORM DI and REM were correlated across all time points. However, this correlation was lost with radiation exposure. From our Principal Component Analysis, ucOC positively interacts with 3-point bending structural parameters and negatively interacts with cortical bone material properties (modulus). In addition, higher levels of ucOC were correlated with better performance in rPVT OPS at later timepoints (90- and 180-days). However, there were no significant correlations found in our Spearman’s correlations across all timepoints. It is not clear if osteocalcin is the osteokine that is responsible for these effects. This seems to indicate that radiation exposure is affecting additional, non-shared pathway(s) between bone and the CNS. Future work could explore additional cytokines to determine what specific pathways are affected.

**Figure 10.**
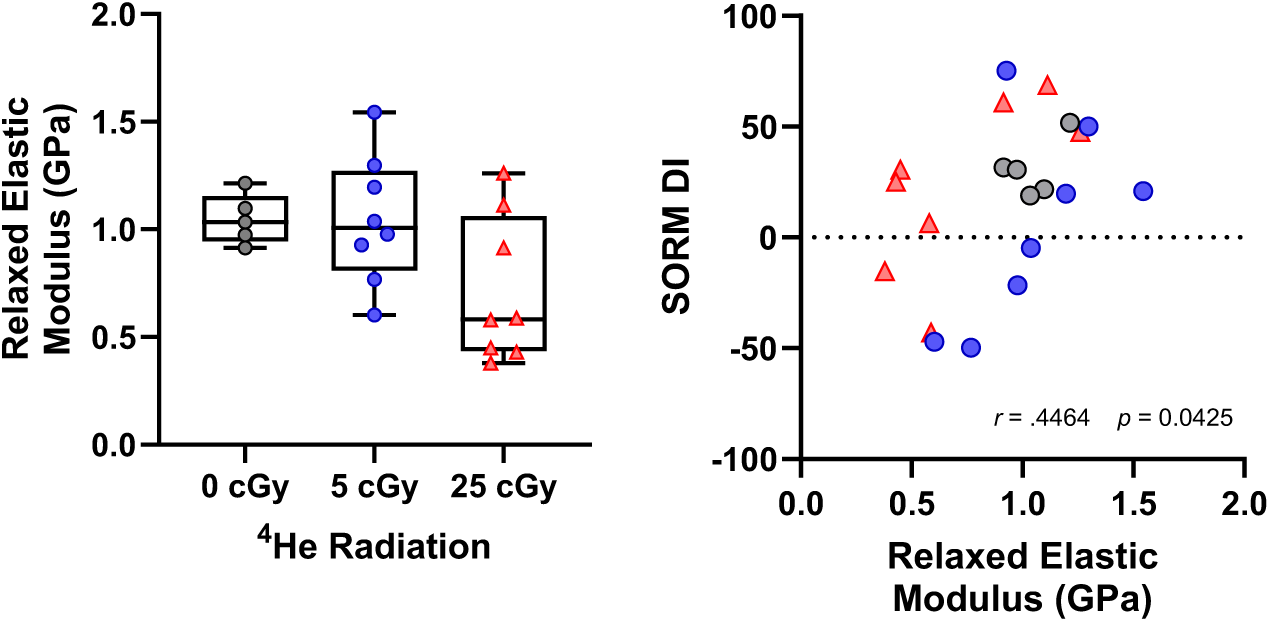
Left: Relaxed Elastic Modulus in rats in the 90-day groups. Right: Significant positive correlation between Relaxed Elastic Modulus (REM) and SORM DI for rats in the 90-day groups.

One limitation of this study is the sample sizes for the bone analyses. The parent study was originally powered for the neurobehavioral evaluation, and the bone analyses were added as a tissue-sharing study. No morphological or histological bone data, or bone activity markers (e.g., Alkaline Phosophtase, TRAP 5B) were included. The biomechanical tests conducted are not able to capture some of the small microstructural changes in the bone that might result from low dose radiation exposure (i.e., introduction of very small pores in the cortical bone). Future work can incorporate other experimental techniques to quantify very small length scale changes (∼1µm) such as coaxial light microscopy, as well as changing the length scale of the spherical indentation methods (i.e., nano vs microindentaion).

In summary, our findings demonstrate that exposure to low doses of ^4^He ions, a significant component of GCR, leads to long-term and progressive deficits in sustained attention and social recognition memory, with no evidence of recovery up to six months post-exposure.

These cognitive impairments are concerning for long-duration space missions, particularly given the potential for exacerbation by other stressors like sleep disruption and spaceflight. While bone mechanical properties were largely unaffected at these low doses, specific parameters showed alterations, and age-related changes likely played a role. Furthermore, we identified circulating cytokines and ucOC as potential biomarkers for radiation-induced neurobehavioral effects, suggesting avenues for future diagnostic development. The sustained nature of these deficits underscores the critical need for effective countermeasures to protect astronaut health and performance during exploration-class missions.

## Acknowledgements

We would like to thank Dr. Robert Hienz for discussions related to design and completion of this study, and Ray Smith, Blanca Bravo, Stacey Perry, Sabrina Lindemon, and Brachman Herzig for their technical assistance. We would also like to thank Dr. Peter Guida, Dr. Adam Rusek, the NASA Space Radiation Laboratory support staff, and the Brookhaven Laboratory Animal Facility technical staff for their help executing all aspects of these studies at Brookhaven National Laboratory. This work was supported by NASA grants 80NSSC22K0022 (CMD) and 80NSSC21K1506 (AGL), USUHS New Faculty Hiring Funds (CMD), and The College of New Jersey MUSE program (AGL), and the NJ Space Grant Consortium (AGL). Some of the authors are employees of the U.S. Government, and this work was prepared as part of their official duties. Title 17 U.S.C. §105 provides that ‘Copyright protection under this title is not available for any work of the United States Government.’ Title 17 U.S.C §101 defines a U.S. Government work as a work prepared by a military service member or employees of the U.S. Government as part of that person’s official duties. The opinions and assertions expressed herein are those of the author(s) and do not reflect the official policy or position of the Uniformed Services University of the Health Sciences, the Armed Forces Radiobiology Research Institute, Department of the Navy, the Department of War, or the US Federal Government. Data from these experiments will be freely available (after an embargo period) through the NASA Open Science Data Repository (OSDR) under study number OSD-892.

